# Live-cell imaging under centrifugation characterized the cellular force for nuclear centration in the *Caenorhabditis elegans* embryo

**DOI:** 10.1101/2024.01.03.574024

**Authors:** Makoto Goda, Michael Shribak, Zenki Ikeda, Naobumi Okada, Tomomi Tani, Gohta Goshima, Rudolf Oldenbourg, Akatsuki Kimura

**Affiliations:** Marine Biological Laboratory, Woods Hole, Massachusetts 02543, USA; Hamamatsu University School of Medicine, Hamamatsu 431-3192, Japan; Nagoya University, Nagoya 464-8602, Japan; National Institute of Genetics, Mishima 411-8540, Japan; Sokendai (Graduate University for Advanced Studies) Mishima, Mishima 411-8540, Japan; Independent Researcher; Biomedical Research Institute, National Institute of Advanced Industrial Science and Technology, Ikeda 563-8577, Japan

## Abstract

Organelles in cells are appropriately positioned, despite crowding in the cytoplasm. However, our understanding of the force required to move large organelles, such as the nucleus, inside the cytoplasm is limited, in part owing to a lack of accurate methods for measurement. We devised a novel method to apply forces to the nucleus of living, wild-type *Caenorhabditis elegans* embryos to measure the force generated inside the cell. We utilized a centrifuge polarizing microscope (CPM) to apply centrifugal force and orientation-independent differential interference contrast (OI-DIC) microscopy to characterize the mass density of the nucleus and cytoplasm. The cellular forces moving the nucleus toward the cell center increased linearly at ∼14 pN/μm depending on the distance from the center. The frictional coefficient was ∼1,100 pN s/μm. The measured values were smaller than previously reported estimates for sea urchin embryos. The forces were consistent with the centrosome-organelle mutual pulling model for nuclear centration. Frictional coefficient was reduced when microtubules were shorter or detached from nuclei in mutant embryos, demonstrating the contribution of astral microtubules. Finally, the frictional coefficient was higher than a theoretical estimate, indicating the contribution of uncharacterized properties of the cytoplasm.

## Introduction

The cell interior is highly crowded^1^. Our understanding of the amount of force required to move a large object inside the crowded cell is limited^2^ and the molecular mechanisms that produces the force are unclear. The nucleus is the largest organelle in animal cells. In some cases, such as pronuclear migration after fertilization, the nucleus moves a long distance^3^. Upon fertilization, the sperm-derived pronucleus forms at a peripheral position where the sperm entered the oocyte. Its migration to the center of the fertilized egg is mainly driven by microtubule asters^4–6^. Microtubules can generate force by dynamically elongating and shrinking or by acting as a rail for molecular motors^7^. They often extend radially from the centrosome, a microtubule organizing center (MTOC), forming an aster. Since the microtubule aster itself is a large cellular structure, the migration of the pronucleus together with the asters in the crowded cytoplasm is expected to require large forces^8^.

The forces generated in the cell to move the pronucleus during its migration were first characterized in fertilized sea urchin eggs^9^. Tanimoto et al. used magnetic tweezers to measure the force required to move the complex consisting of the microtubule asters (extending up to ∼50 μm to reach the cell cortex), sperm-derived pronucleus, and oocyte-derived pronucleus. They revealed that the microtubule aster produces a force of 580 ± 21 pN to move the nuclei-aster complex, whose frictional coefficient is 8,400 ± 280 pN s/μm. Pulling of the microtubules and thus the nuclei-aster complex at the cytoplasm by cytoplasmic dynein is thought to generate the migration force in sea urchin^9–11^ and in other organisms^6,12,13^. However, another report argues that dynein is not required for the pronuclear migration in sea urchin^14^. Thus, the molecular mechanism underlying force production and the high frictional coefficient of the nuclei-aster complex remain unclear in sea urchin eggs.

The nematode *Caenorhabditis elegans* is another popular model owing to the ease of live imaging and gene manipulation. The forces generated by the microtubule aster and the frictional coefficient to move the nuclei-aster complex for the pronuclear centration have not been measured experimentally in this system. In theory, the frictional coefficient can be estimated as follows. The frictional coefficient, *F*_drag_/*V*, of a spherical object inside a simple viscous fluid (i.e., Newtonian fluid) follows Stokes’ law^15^, as *F*_drag_/*V* = 6π*ηR*, where *F*_drag_ is the force for dragging the sphere, *V* is the velocity of the sphere movement, *η* is the viscosity of the medium, and *R* is the radius of the sphere. In *C. elegans*, the sperm-derived pronucleus is a spherical object with a radius of 4.5 μm, with an estimate of 0.2 pN s/μm^2^ for the viscosity of the cytoplasm^16^. Applying these values to Stokes’ equation, the frictional coefficient is ∼20 pN s/μm. The movement of the nucleus has a compressive effect on the cytoplasm, and simulation studies have shown that the effect of confinement increases the coefficient by 3.3-fold^8,17^. Furthermore, the nuclei-aster complex is not a smooth sphere; a simulation study compares it to a porous medium with a 6-fold increased frictional coefficient^8^.

This calculation is consistent with an experimental observation that the movement of a sperm pronucleus with microtubule asters is 4.4-fold slower than that of an oocyte pronucleus without microtubule asters^18^. In sum, the theoretically estimated value of the frictional coefficient of the nuclei-aster complex in the *C. elegans* embryo is ∼300 pN s/μm, about 30-fold smaller than the experimentally measured value in sea urchin (8,400 pN s/μm)^9^. The maximum speed of pronuclear migration in *C. elegans* is ∼0.1 μm/s. The force required to generate such speed with the frictional coefficient of 300 pN s/μm is estimated to be 30 pN, about 20-fold smaller than the experimental measurement in sea urchin (580 pN). To resolve these discrepancies, experimental measurements of force for *C. elegans* pronuclear migration are needed.

Forces require for positioning the mitotic spindle in *C. elegans* were measured by Garzon-Coral et al.^16^ In contrast, the measurement of forces related to nuclear positioning in the *C. elegans* embryo has not been reported. The centrifuge microscope^19^ is a promising tool to measure intracellular forces, as it allows live imaging of a microscopic specimen under centrifugal forces. If the densities of the nucleus and cytoplasm are different, the nucleus should move depending on the centrifugal speed according to the following formula: *F*_cfg_ = *Δρ*×*NV*×RCF×*g*. *F*_cfg_ is the centrifugal force acting on the nucleus, *Δρ* is the density difference between the nucleus and the cytoplasm, *NV* is the volume of the nucleus, RCF is the relative centrifugal force, and *g* is the gravitational acceleration (9.8 m/s^2^). Even if the density difference is small, the nucleus will be moved by the centrifugal force if the rotational speed (RCF) is high enough. The centrifuge polarizing microscope (CPM) invented by Inoué and colleagues is capable of imaging under a rotational speed that generates up to ∼10,000 ×*g*, the fastest to date, to the best of our knowledge^19,20^. The CPM has been used to apply forces to various biological samples^21–23^. In this study, by applying centrifugal forces using the CPM, we measured the cellular forces and frictional coefficient for nuclear centration in *C. elegans* embryos.

## Results

### Movement of the nucleus in the *C. elegans* embryo under centrifugal forces

In the CPM^19^, the rotor spins between the objective (40×, NA 0.55) and condenser lenses (Fig. 1A). The specimen in the rotor is illuminated stroboscopically by 6-ns laser pulses, which are synchronized to the exact timing when the specimen comes under the objective lens. Therefore, despite the fast rotation of the rotor, the camera image is stationary with high resolution, up to 1 μm.

**Figure 1.**
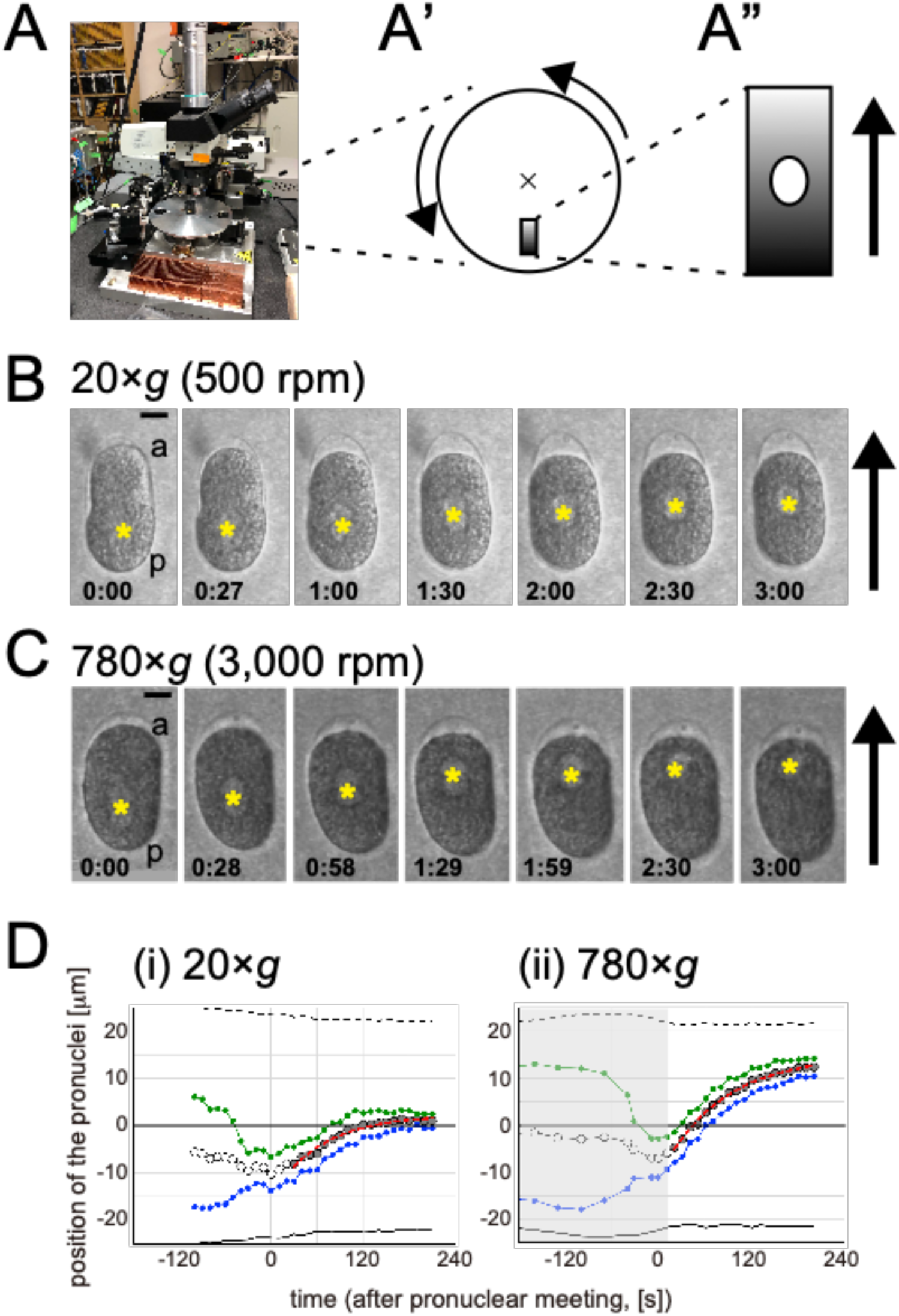
Observation of *C. elegans* embryos using a centrifuge polarizing microscope (CPM). (A) View of the CPM. (A’, A”) Schematic drawings of the top views of the rotation stage (circle) and sample chamber (rectangle). The direction of the rotation of the stage is shown with arrows in (A’). The direction of the center of the rotation (the cross in A’) is shown with arrow in (A”). The gradient of the grey color indicates the density gradient of the Percoll. The ellipsoid in (A”) indicates the *C. elegans* embryo. (B, C) Time-lapse images of a *C. elegans* embryo at the 1-cell stage obtained using the CPM with the indicated rotation speed. Asterisks indicate the position of the pronuclei. a: anterior pole, p: posterior pole. The arrows indicate the center of the rotation. Time is indicated in min:sec. Time 0:00 was set to the time when the pronuclei met. (B) 20 ×*g* (500 rpm). (C) 780 ×*g* (3,000 rpm). Scale Bar, 10 μm. (D) Representative plots of the position of the nuclei along the long (anterior-posterior) axis of the cell. (i) 20 ×*g* (500 rpm), (ii) 780 ×*g* (3,000 rpm). Position = 0 [μm] is at the cell center. The position is negative for the posterior half of the cell. Time = 0: pronuclear meeting. The dotted and solid line indicate the positions of the anterior and posterior pole of the cell. The green and blue circles indicate the positions of the oocyte-, and sperm-derived pronucleus, respectively. The white and gray circles indicate the center of the two pronuclei. The grey circles indicate the data used for the fitting. The red dotted line indicates the best-fit curve based on the equation *x* = *L*_eq_ × [1 - exp{-(*t* - *t*_0_)/*τ*}] (see Methods). In (C) and (D-ii), the rotation speed was lower than 3,000 rpm (780 ×*g*) until +19 s (grey region in (D-ii)). Specifically, the rotation speed was 500 rpm (20 ×*g*) until just before the meeting (-31 s) to ensure pronuclear meeting, and increased to 3,000 rpm. It took ∼50 s for the speed to reach 3,000 rpm.

We mounted the *C. elegans* embryo in a chamber designed for the CPM. Inside the chamber, a density gradient of Percoll was established and the position of a floating embryo was fixed in the chamber relative to the Percoll gradient (Fig. 1A’, 1A”). The anterior pole of the *C. elegans* embryo always faced the center of the point of rotation during the one-cell stage (Fig. 1B, 1C). The reproducible orientation of the anterior pole to the center was caused by the anterior localization of a low-mass-density material (see the section after the next one).

We then observed the movement of the pronuclei. Without centrifugation (for example, see Fig. 1 in Kimura and Onami^6^), the sperm- and oocyte-derived pronuclei formed at posterior and anterior positions, respectively, and met and attached to each other at a posterior position (pronuclear meeting, time point 0 in this study). Following attachment, the pronuclei moved together toward the center of the embryo, where they were stabilized until mitosis. Under a low centrifugal speed (500 rpm, 20 ×*g*), the movement of the pronuclei was similar to that in the non-centrifugation condition (Fig. 1B, Supplemental Movie S1). In contrast, when we increased the centrifugal speed to 3,000 rpm (780 ×*g*), after the pronuclear meeting, the pronuclei passed the center of the cell and moving further towards the center of the rotation stage (Fig. 1C, Supplemental Movie S2). These observations revealed that (i) the pronuclei can be moved by centrifugal forces produced by the CPM and (ii) the mass density of the pronuclei is lower than that of the cytoplasm.

Even though nuclei migrated to the anterior pole, the mitotic spindle, which is formed after nuclear envelope breakdown, moved back to the normal position and divided normally (Supplemental Movies S1 and S2). This is likely because the mitotic spindle is not lighter than the cytoplasm and the centrifugal force does not act on it.

Embryos subjected to 780 ×*g* during pronuclear migration and the first cell division hatched normally after unmounting from the CPM chamber and culturing under normal gravity. We concluded that centrifugal force of up to 780 ×*g* is a mild perturbation and does not cause detectable damage to cellular structures related to pronuclear migration. We thus used this experimental setup to relate the external force provided by centrifugation and the movement of the pronuclei, to quantitatively evaluate the forces produced by the cell.

### Toward the calculation of the centrifugal force: nuclear volume

The centrifugal forces acting on an object can be calculated as *F*_cfg_ = *Δρ*×*NV*×RCF×*g*. *Δρ* is the density difference between the object and the suspending medium, *NV* is the volume of the object, RCF is the relative centrifugal force, and *g* is the gravitational acceleration (9.8 m/s^2^). To calculate the force from RCF, which is a function of the centrifuge speed, the parameters *NV* and *Δρ* are necessary.

We quantified the nuclear volume (*NV*) from confocal microscopy images of cells whose nuclear membrane was fluorescently labeled (Table 1 and Supplemental Fig. S1). We focused on the movement of the pronuclei after the sperm- and oocyte-derived pronuclei meet. This is because high centrifugal force applied before the two pronuclei meet can prevent the meeting, making the comparison between different forces difficult. The total volumes of the two pronuclei were 731 ± 93 μm^3^ (mean ± s.d. from 13 embryos at 51 timepoints).

**Table 1.** Forces and related values measured or calculated in this study.

|  | symbol | unit | mean±s.d. | n |
| --- | --- | --- | --- | --- |
| nuclear volume (SPN+OPN*) | $NV$ | $\mu\text{m}^3$ | 731±93 | 51 |
| density difference between<br>nucleus and cytoplasm | $\Delta\rho$ | mg/mL | 37.4±6.8 | 6 |
| slope: $L_{eq}$ vs RCF* | $\beta_{RL}$ | $\mu\text{m}/[\text{xg}]$ | $(1.88\pm0.26) \times 10^{-2}$ | 33 |
| stiffness-1 | $K_1$ | pN/um | 14.3±3.9 | |
| slope: $V_0$ vs. RCF | $\beta_{RV0}$ | $\mu\text{m}/\text{s}$ | $(2.41\pm0.25) \times 10^{-4}$ | 33 |
| frictional coefficient (SPN+OPN) | $C_{\text{fric}}$ | pN s/$\mu\text{m}$ | 1,110±288 | |
| stiffness-to-friction ratio | $R_{KC}$ | /s | $(2.64\pm1.38) \times 10^{-2}$ | 10 |
| stiffness-2 | $K_2$ | pN/um | 29.3±17.1 | |
| slope: $V_0$ vs RCF (N2, SPN) | $\beta_{RV0\_WT\_S}$ | $\mu\text{m}/\text{s}$ | $(2.08\pm0.66) \times 10^{-4}$ | 5 |
| slope: $V_0$ vs RCF (zyg9, SPN) | $\beta_{RV0\_z9\_S}$ | $\mu\text{m}/\text{s}$ | $(4.28\pm0.43) \times 10^{-4}$ | 8 |
| slope: $V_0$ vs RCF (zyg12, SPN) | $\beta_{RV0\_z12\_S}$ | $\mu\text{m}/\text{s}$ | $(3.28\pm0.37) \times 10^{-4}$ | 8 |
| frictional coefficient (N2, SPN) | $C_{\text{fric\_WT\_S}}$ | pN s/$\mu\text{m}$ | 644±255 | |
| frictional coefficient (zyg-9, SPN) | $C_{\text{fric\_z9\_S}}$ | pN s/$\mu\text{m}$ | 313±81 | |
| frictional coefficient (zyg-12,<br>SPN) | $C_{\text{fric\_z12\_S}}$ | pN s/$\mu\text{m}$ | 409±11 | |

\* SPN = sperm-derived pronucleus, OPN = oocyte-derived pronucleus, RCF = relative centrifugal force

### Toward the calculation of the centrifugal force: mass density

To quantify *Δρ*, which is the density difference between the pronuclei and the cytoplasm, we utilized orientation-independent differential interference contrast (OI-DIC) microscopy^24,25^. OI-DIC measures the optical path difference (OPD) within a microscope image. From the OPD and the thickness of the object, the refractive index can be calculated. Because the refractive index is proportional to the dry mass and because the refractive increments of most substances in cells are approximately the same^26–28^, mass density in the living cell can be calculated from the refractive index^29^.

When we imaged the *C. elegans* embryo with OI-DIC, the refractive index outside the cell but inside the eggshell was comparable to that of the buffer and apparently lower than that of the cell (Fig. 2A (iii), asterisk). This area is referred to as the EEM (extra embryonic matrix)^30^. The low refractive index of the EEM concentrated at the anterior side of the embryo suggested that the mass density of the anterior side is lower than that of the posterior side. The mechanism by which the EEM localizes to the anterior side of the embryo will be published elsewhere (ZI, AK et al., in preparation). This observation is consistent with our CPM observation that the anterior pole always faces toward the center of the rotation (Fig. 1B, 1C). These results further indicated that OI-DIC is a reliable method to estimate the mass density inside the cell.

**Figure 2.**
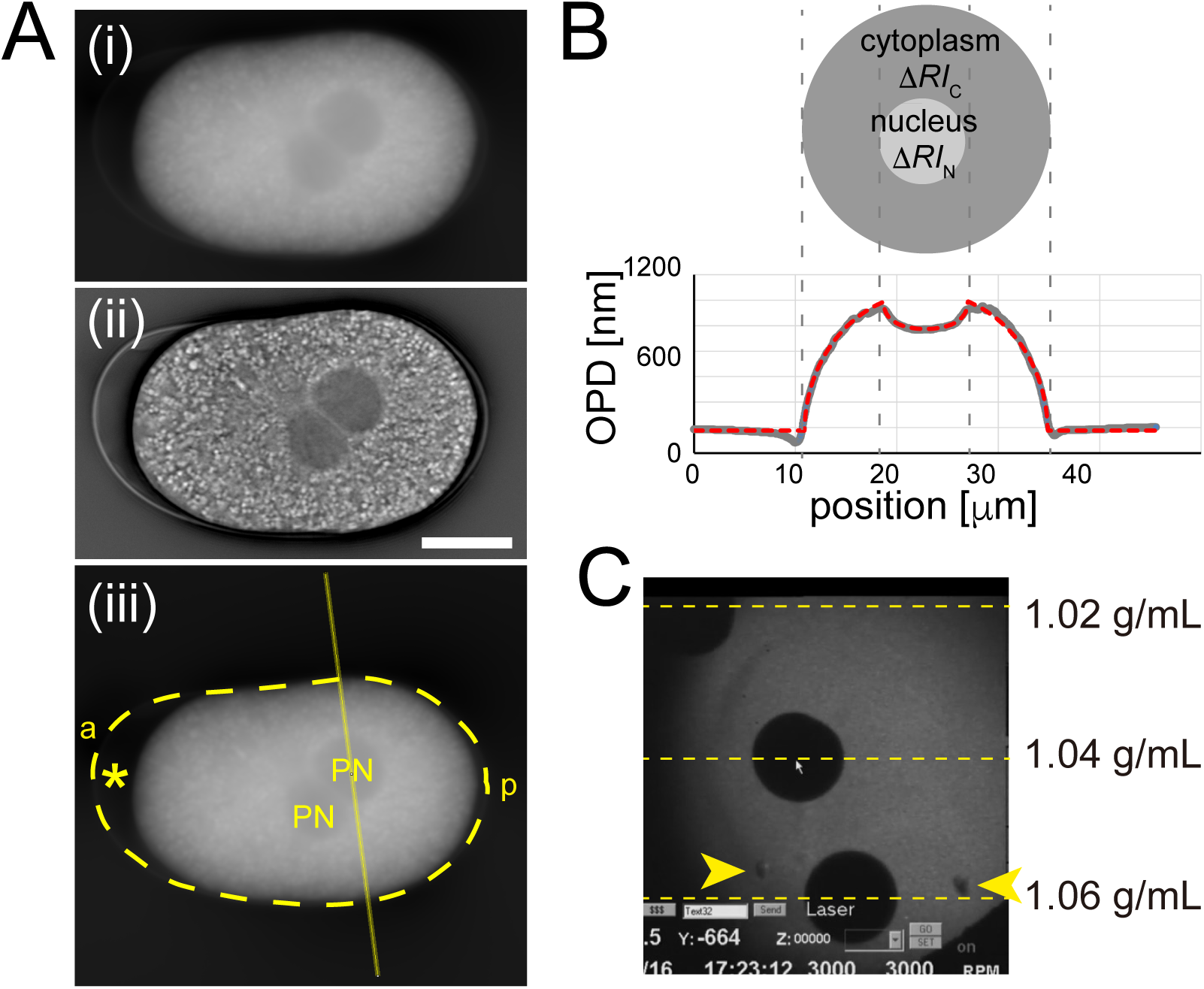
Characterization of mass density using OI-DIC. (A) OI-DIC image of the *C. elegans* embryo at the pronuclear meeting stage. (A-i) Values of OPD (optical path difference) are visualized by the degree of whiteness. (A-ii) Image of the embryo processed using the inverse Riesz transform to visualize the pronuclei and eggshell. (A-iii) The same image shown in (i) but with a line passing the center of a pronucleus drawn to quantify the OPD in (B) and the position of the eggshell determined from (ii) visualized as a dotted curve. *, extra embryonic matrix (EEM). PN, the two pronuclei. a: anterior pole, p: posterior pole. Scale bar, 10 μm. (B) Example of a line scan of the OPD perpendicular to the long axis of the embryo (grey) and the result of fitting the refractive indices of the cytoplasm and nucleus (red dotted line) using expressions given in the Methods section. The upper schematic shows the cross section of the embryo (yellow straight line in (A-iii)). The OPD will be [baseline] + *ΔRI*_C_ × *T*_C_ + *ΔRI*_N_ × *T*_N_, where *ΔRI*_C_ and *ΔRI*_N_ are the differences in refractive index (RI) against the buffer (water) for the cytoplasm and the nucleus, and *T*_C_ and *T*_N_ are the thickness of the cytoplasm and nucleus, respectively (see Methods). (C) Estimation of the mass density of the embryo by comparing the position of the embryos and beads with the known mass density in the CPM. Yellow-dashed lines indicate the position of the beads with the indicated density. Yellow arrowheads indicate the positions of the embryos in the chamber.

The lower refractive index of the nuclear region than the cytoplasm (Fig. 2A) indicated that the mass density of the nucleus is lower than that of the cytoplasm, in agreement with the movement of the nucleus depending on the centrifugal force (Fig. 1). From the 2-dimensional map of the OPD (Fig. 2A (i)), we quantified the densities of the cytoplasm and nucleus by fitting the OI-DIC data to a formula assuming that the cross section of the nucleus and the cell perpendicular to the long axis of the *C. elegans* embryo are circular (Fig. 2B, see Methods). As calculated from the OPD, thickness, and refractive index of the buffer (1.33), the density of the cytoplasm was 1,063 ± 7.8 mg/mL and the density difference between the nucleus and the cytoplasm (*Δρ*) was 37.4 ± 6.8 mg/mL (mean ± s.d. from 6 nuclei of 5 embryos).

The conversion of the OPD to the density difference of cell compartments assumed a linear relationship between the refractive index and dry mass, and a specific refractive index increment of 0.0018 [100 mL/g]^27,29^. To validate this assumption, we measured the mass density of the embryo with an independent method, using the CPM. We mounted the embryo together with colored standard beads of known density. As we applied centrifugal force to the mixture, the positions of the standard beads separated along the Percoll density gradient. The positions of embryos were almost identical to that of a bead with a known density of 1.06 g/mL (Fig. 2C), which agreed with the mass density of the cytoplasm (1,063 ± 7.8 mg/mL) estimated using OI-DIC. In conclusion, we adopted a density difference (*Δρ*) of 37 mg/mL to calculate the centrifugal force on pronuclei.

### Strategy to quantify cellular forces for nuclear centration and the frictional coefficient of nuclei-aster complex

From the abovementioned measurements of nuclear volume and density difference between the nucleus and cytoplasm, we can calculate the amount of centrifugal force applied to the nuclei (*F*_cfg_). The purpose of this study is to quantify the force produced by the cell to move and maintain the nuclei at the center of the cell (*F*_centration_). In addition, the drag force (*F*_drag_) works to move nuclei in the opposite direction. We assume these three forces are the major forces acting on the pronuclei and are balanced as follows: *F*_centration_ + *F*_drag_ = *F*_cfg_ (Fig. 3A (i)). The assumption is reasonable because, without centrifugal force (*F*_cfg_ = 0), the nuclei stop (*F*_drag_ = 0) at the center of the cell, where *F*_centration_ should be 0.

**Figure 3.**
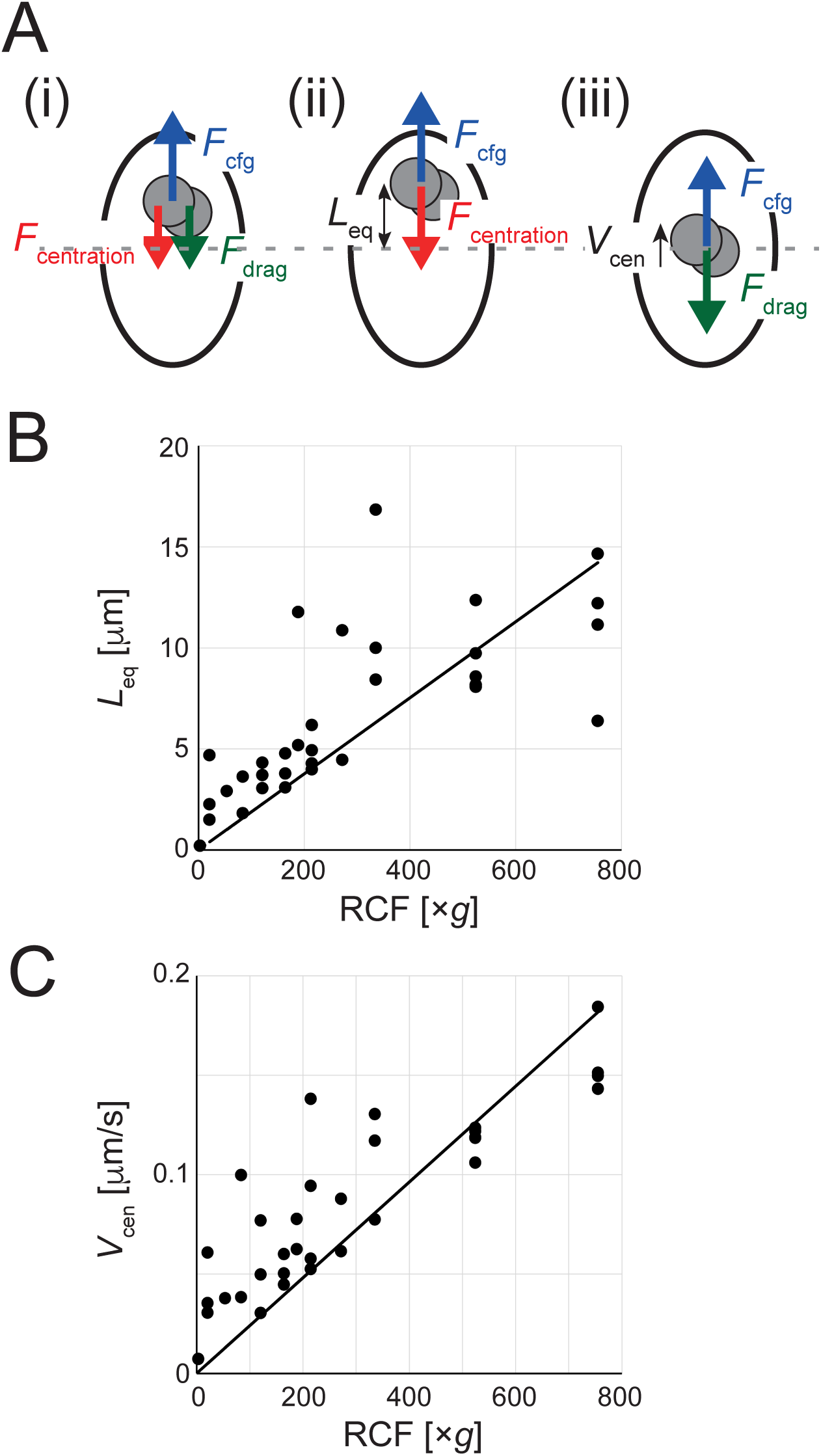
Quantification of the stiffness (*K*_1_) and the frictional coefficient (*C*_fric_) from the movement of the nucleus under the centrifugation. (A) Scheme of the force balance on the nuclei (grey circles) inside the cell (ellipsoid) under centrifugation. (A-i) The centrifugal force (*F*_cfg_, blue) is applied to the nuclei because they are lighter than the cytoplasm. The cellular force (*F*_centration_, red) acts on the nucleus to bring them to the cell center (grey dotted line). When the nuclei moves (toward the upper in this example), the drag force (*F*_drag_, green) acts toward the opposite direction. These three forces should be balanced. (A-ii) When the nuclei stop moving, *F*_drag_ will be zero, and *F*_cfg_ and *F*_centration_ will be balanced. (A-iii) When the nuclei is at the center, *F*_centration_ will be zero, and *F*_cfg_ and *F*_drag_ will be balanced. (B) Relationship between the displacement of the pronuclei from the center of the cell when the pronuclei stop moving (*L*_eq_) against the centrifuge speed (RCF, ×*g*). Each point represents one experiment. The line is the linear regression line that crosses the origin: *L*_eq_/RCF = (1.88 ± 0.26)×10^-2^ [μm/×*g*]. (C) Relationship between the velocity of the pronuclei passing the center of the cell (*V*) against the centrifuge speed (RCF, ×*g*). Each point represents one experiment. The line is the linear regression line that crosses the origin: *V*/RCF = (2.41 ± 0.25)×10^-4^ [μm/s×*g*].

Using this equation, we quantified *F*_centration_ and *F*_drag_. When the nuclei are not moving, the drag force is zero (*F*_drag_ = 0). In this condition, the centration force equals the centrifugation force (*F*_centration_ = *F*_cfg_) (Fig. 3A (ii)). Similarly, when the nuclei are at the center, the centration force should be zero (*F*_centration_ = 0). In this condition, the drag force equals the centrifugation force (*F*_drag_ = *F*_cfg_) (Fig. 3A (iii)). We therefore focused on the position where the nuclei stop moving to determine *F*_centration_ and the speed of nuclei when they pass the center of the cell to determine *F*_drag_ under various centrifugation conditions.

### *F*_centration_: the force produced by the cell to move and maintain the nuclei at the cell center

We tracked the positions of pronuclei under various centrifugal speeds (Fig. 1D) and fitted the position of the center of the pronuclei, *x*, as a function of time, *t*, with a formula

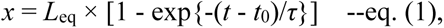

where *L*_eq_ is the position where the pronuclei stop moving (i.e., *F*_cfg_ = *F*_centration_), *t*_0_ is the time when the pronuclei pass the center, and *τ* is a characteristic time-scale for the movement (see Methods). We detected a roughly linear relationship between the centrifugal force and *L*_eq_, i.e., how far the pronuclei were displaced from the center (Fig. 3B). This indicates that the cellular machinery to bring nuclei to the center of the cell behaves like a Hookean spring: the further the nuclei are displaced (*x*), the stronger the forces generated towards the center (*F*_centration_ = - *K* × *x*, where *K* is the stiffness of the centering spring). We concluded that the Hookean spring mechanism accounts for the majority of the relationship between force and distance. However, the imperfect fit to the linear line (the coefficient of determination, R^2^ = 0.81) suggested that additional mechanisms contribute, such as the confinement effect^8^. The slope of the fitted regression line was (1.88 ± 0.26) × 10^-2^ μm/[×*g*] (mean ± s.d., *n* = 33) (Fig. 3B). Based on the nuclear volume and density difference measured in this study, the stiffness *K*_1_ of the ‘Hookean spring’ for the centration of the nuclei in the cell was *K*_1_ = 14.3 ± 3.9 pN/μm (mean ± s.d.) (Table 1). This is the first direct measurement of the centration force of nuclei in *C. elegans* embryos, and the first measurement of the nuclear centration force utilizing centrifugal forces.

### *F*_drag_: drag force

To characterize the drag force for nuclear centration, we focused on the speed of the nuclei when they pass the center, *V*_cen_ (i.e., *F*_centration_ = 0). The speed was calculated from the fitting to eq. (1) (red dotted line,in Fig. 1D) as *V*_cen_ = *L*_eq_/*τ*. We plotted the speed against centrifugal force in RCF (Fig. 3C), revealing roughly a linear relationship. In viscosity-dominant conditions, or a low Reynolds number regime^31^, the drag force (*F*_drag_) is proportional to the velocity of an object (*V*), and the frictional coefficient *C*_fric_ is the ratio between them, *C*_fric_ = *F*_drag_/*V*. The intracellular environment is considered to be viscosity-dominant, and our measurements agree with this notion. We concluded that the proportional relationship largely explains the relationship between force and velocity. However, the imperfect fit to the linear line (the coefficient of determination, R^2^ = 0.88) suggested that additional mechanisms contribute, such as the viscoelasticity of the cytoplasm^16^ or the non-spherical nature of the nucleus-aster complex^8^. The slope of the regression line was (2.41 ± 0.25) × 10^-4^ μm/s ×*g* (mean ± s.d., *n* = 33). Based on the values of nuclear volume and density difference, the frictional coefficient, *C*_fric_, was calculated to be 1,110 ± 288 pN s/μm (mean ± s.d.). This value is higher than a theoretical estimation^8,18^, which was 300 pN s/μm for the sperm-derived pronucleus-aster complex (see Introduction). In the present experiment, we examined the movement of the complex consisting of the fused sperm- and oocyte-derived pronuclei and aster.

Because the size of the aster is the major determinant of the frictional coefficient^8^ and the size does not change substantially before and after the fusion of the two pronuclei^6^, the frictional coefficient for one nucleus is estimated to be similar to two nuclei (∼1,110) or at least its half (∼560 pN s/μm). Therefore, our measurement suggests that uncharacterized components affect the viscosity of the cytoplasm (see Discussion).

### Calculating the centration force from the frictional coefficient and the migration without centrifugation

In this section, we estimate the stiffness of the centration force generated by the cell,

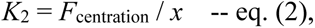

in an independent manner from our previous estimation (Fig. 3B, *K*_1_ = 14.3 ± 3.9 pN/μm). Here, *x* is the position of the center of the pronuclei from the cell center. In the following independent estimation, we use the centration movement of the nucleus under normal (no centrifugation) conditions. In this condition, *F*_centration_ = *F*_drag_ because *F*_cfg_ = 0. From the approximately linear relationship between *F*_drag_ and nuclear velocity, *V*, in Fig. 3C:

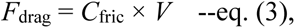

where *C*_fric_ = 1,110 ± 288 pN s/μm. The eqs. (2) and (3) provide the differential equation

*V* = d*x*/d*t* = - (*K*_2_/*C*_fric_) × *x* and thus

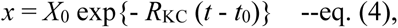

where, *R*_KC_ = *K*_2_/*C*_fric_ describes the ratio of *K*_2_ to *C*_fric_, and *X*_0_ is the position, *x*, at *t* = *t*_0_. We tracked the movement of the pronuclei under normal conditions (Fig. 4A).

**Figure 4.**
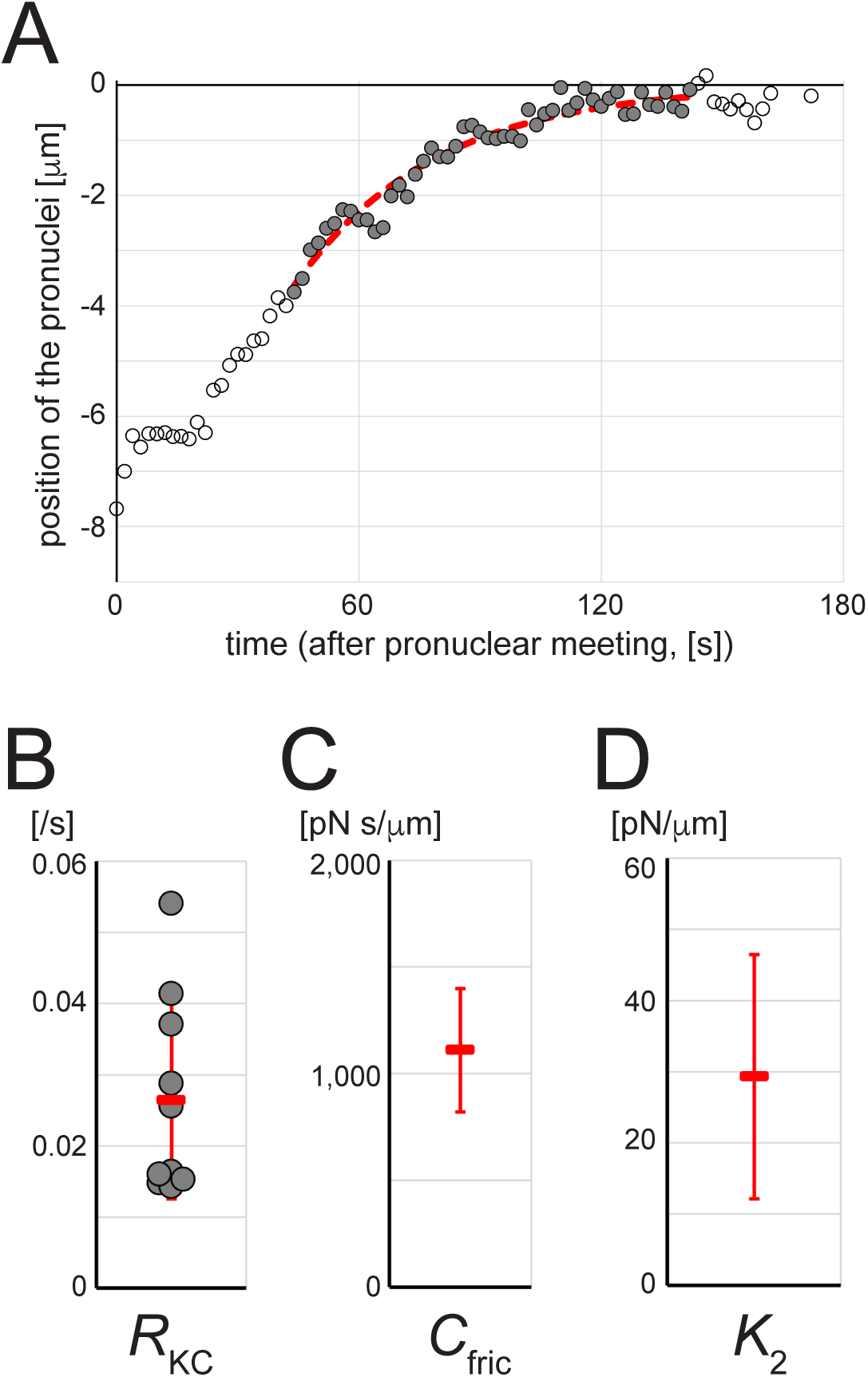
Independent quantification of the stiffness (*K*_2_) from the pronuclear migration in non-centrifuge conditions. (A) Representative plot of the position of the center of the two pronuclei along the long (anterior-posterior) axis of the cell. Position = 0 [μm] is at the cell center. The position is negative for the posterior half. Time = 0: pronuclear meeting. The filled circles indicate the data used for the fitting (see Methods. The red dotted line indicates the best-fit curve based on the equation *x* = *X*_0_ exp{- *R*_KC_ (*t* - *t*_0_)}. (B) The *R*_KC_ (= *K*_2_/*C*_fric_) values obtained by the fitting in Fig. 4A, from 10 embryos (circles). The mean and s.d. are indicated with red. (C) The mean and s.d. of the *C*_fric_ calculated in Fig. 3C. (D) The mean and s.d. of the *K*_2_ calculated from the values in Fig. 4B and 4C, the relationship *K*_2_ = *R*_KC_ × *C*_fric_, and the law of error propagation.

We fitted the movement trajectories to eq. (4) (Fig. 4A, dotted line) and obtained *R*_KC_ = *K*_2_/*C*_fric_ = (2.64 ± 1.38) × 10^-2^ /s (mean ± s.d., *n*=10) (Fig. 4B). Using the frictional coefficient of *C*_fric_ = 1,110 ± 288 pN s/μm (Fig. 3C, Fig. 4C) and the law of error propagation, we obtain the stiffness of the centering machinery as *K*_2_ = 29.3 ± 17.1 pN/μm (mean ± s.d.) (Fig. 4D, Table 1). This value is comparable to the abovementioned, independent estimate of the stiffness, *K*_1_ = 14.3 ± 3.9 pN/μm (Fig. 3B, Table 1). The similarity between the stiffness values obtained under centrifugal and non-centrifugal conditions supports the notion that the force production by the centration machinery in not affected by our centrifugation procedure. The larger standard deviation of *K*_2_ compared with that of *K*_1_ led us to conclude that *K*_1_ (14.3 ± 3.9 pN/μm) is more reliable than *K*_2_ as the stiffness of the centration mechanism.

### Measurement of nuclear motion in mutant cells

The force to center the pronuclei has been quantified in sea urchin eggs using magnetic tweezers^9^. The present study, quantifying the force in *C. elegans,* not only adds another organism for comparison but also provides a way to characterize gene functions for nuclear centration, as we can easily manipulate gene activity in *C. elegans*. As a proof of concept, we examined the nuclear movement of *zyg-9* and *zyg-12* mutant embryos under the centrifugal force (Fig. 5). *zyg-9* encodes a XMAP215 (Xenopus Microtubule-associated protein) homologue, which promotes the elongation of microtubules^32–34^ (Fig. 5A). The *zyg-12* gene encodes a functional KASH protein in *C. elegans* essential for the attachment of microtubule asters and both sperm and oocyte pronuclei^35^ (Fig. 5A). In *zyg-9* or *zyg-12* temperature sensitive mutant embryos, active pronuclear migration does not occur^32,35,36^ at the restrictive temperature because microtubule growth and attachment to the pronucleus are required^36^.

**Figure 5.**
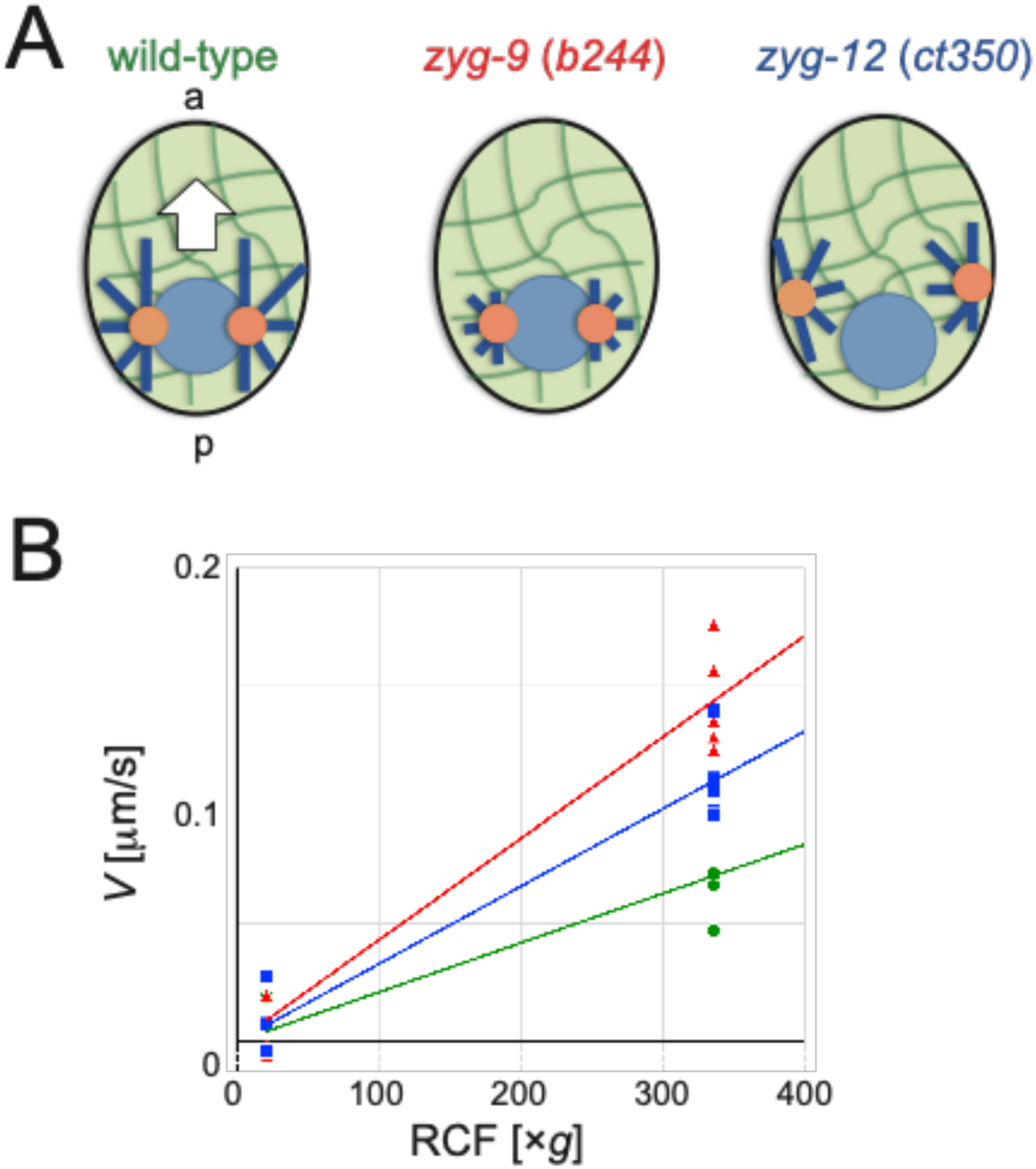
Frictional coefficient in *zyg-9* versus *zyg-12* mutant embryos. (A) Schematic drawings of the microtubule organization in the control (wild-type), *zyg-9*, and *zyg-12* mutant embryos during pronuclear migration. In the control, the sperm-derived pronucleus (pale blue circle) is associated with the centrosome (orange circles) and the microtubule asters (dark blue lines), to move through the cytoplasm (pale green with dark green networks of cytoskeleton and organelles) toward the center (white arrow). In *zyg-9* mutant, microtubule elongation is impaired, and centration is defective. In *zyg-12* mutant, the centrosomes dissociate from the nucleus and the centration of the nucleus is defective. As we focused on the sperm-derived pronucleus in this section, the oocyte-derived pronucleus is not shown in the schematics. a: anterior pole, p: posterior pole. (B) Relationship between the velocity (*V*) of the sperm-derived pronucleus against the centrifuge speed (RCF, ×*g*). (Green circle) Control (wild-type, velocity of the sperm-derived pronucleus when it passes the center of the cell). (Red triangle) *zyg-9* (*b244*ts). (Blue square) *zyg-12* (*ct350*ts). The velocity of the sperm-derived pronucleus in *zyg-9* and *zyg-12* mutants was constant regardless of its position, possibly due to the defect in centration force, and thus only depended on the centrifuge speed. The linear regression line that passes through the origin was drawn. (Green) *V*/RCF = (2.08 ± 0.66)×10^-4^, (red) (4.28 ± 0.43)×10^-4^, and (blue) (3.28 ± 0.37)×10^-4^ [μm/s x*g*].

Under slow centrifugation (500 rpm, 20 ×*g*), the pronuclei showed little movement (Fig. 5B), as observed under no centrifugation. We increased the centrifugation speed for the *zyg-9* (*b244*ts) and *zyg-12* (*ct350*ts) mutant embryos to 2,000 rpm (340 ×*g*) at the timing of the relaxation of the pseudo-cleavage furrow, which corresponds to the timing of pronuclear meeting in control cells. Both the sperm and oocyte pronucleus moved centripetally at almost a constant speed. We measured the speed as a function of rotational speed to calculate the frictional coefficient in *zyg-9* and *zyg-12* mutant embryos. The slopes of the regression line were (4.28 ± 0.43) × 10^-4^ μm/s ×*g* (mean ± s.d., *n*=8) for the *zyg-9* mutant and (3.28 ± 0.37) × 10^-4^ μm/s ×*g* (mean ± s.d., *n*=8) for the *zyg-12* mutant. Note that these values are for one (sperm-derived) pronucleus but not for the two attached pronuclei, as measured for wild-type embryos above. As the wild-type control for the frictional coefficient of the sperm-derived pronucleus, unlike the previous experiments, we increased the centrifugation speed before the pronuclear meeting and measured the velocity of the sperm-derived pronucleus when it crossed the center of the cell. The slope of the regression line was (2.08 ± 0.66) × 10^-4^ μm/s ×*g* (mean ± s.d., *n* = 5) for the sperm-derived pronucleus in the wild-type. Based on the volume of the pronucleus and the density difference, the frictional coefficient for the sperm-derived pronucleus were 313 ± 81, 409 ± 11, and 644 ± 255 pN s/μm (mean ± s.d.) for *zyg-9*, *zyg-12*, and wild-type, respectively. In this measurement, the frictional coefficient of one pronucleus in the wild-type control (∼600 pN s/μm) was about half that of two pronuclei in our previous measurement (∼1,100 pN s/μm, Fig. 3C). The frictional coefficient was reduced further to about half in *zyg-9* and *zyg-12* pronuclei. This was reasonable because the microtubule asters associated with the pronucleus should interfere with the movement inside the cytoplasm^8,18^.

Quantitatively, the difference of about two-fold was smaller than the previous theoretical estimation that the microtubule aster should increase the frictional coefficient 6-fold^8^.

## Discussion

### On the value of the centration force

We established a new experimental setup using a centrifuge microscope to quantify the force required for nuclear migration toward the center of the cell. The force was almost proportional to the distance of the nuclei from the center. The force per displacement required to move the nuclei was 14 ± 4 pN/μm (stiffness, *K*_1_), consistent with an independent estimation based on the frictional coefficient and the velocity of nuclei in non-centrifuge condition (stiffness, *K*_2_). The maximum force produced during migration was about 100 pN, based on the frictional coefficient (∼1,000 pN s/μm) and maximum velocity of the nuclei (0.1 μm/s, Fig. 4A).

The magnitude of the force required to displace the nuclei at prophase of the one-cell stage measured with the CPM in this study (14 pN/μm) was very similar to that required to displace a pole of the mitotic spindle at metaphase measured using magnetic tweezers (16 pN/μm)^16^, despite the differences in cell cycle (prophase vs. metaphase) and cargos (nuclei vs. spindle). This similarity suggests that the centering of the pronuclei at prophase and the mitotic spindle at metaphase are accomplished by a common mechanism, although the mechanisms proposed in past studies differ^6,12,16,37^.

The maximum force produced for *C. elegans* pronuclear migration was about 1/6 of the force measured in a sea urchin (580 pN)^9^. A simple explanation for this discrepancy is the difference in cell size (an ellipsoid with a long axis of 50 μm and short axes of 30 μm for the *C. elegans* embryo, and a sphere with a diameter of 90 μm for the sea urchin egg). If the driving forces are produced throughout the cytoplasm and thus are dependent on the length of the microtubule, as proposed in these organisms^6,9–12,38^, it is reasonable for the generated force to depend on the cell size. However, the force-producing mechanism for centration is currently under debate and may involve microtubule pushing against the cell cortex^14,16^ or microtubule pulling from the cell cortex^37,39^.

### On the value of frictional coefficient of the nuclei-aster complex

The frictional coefficient of the nuclei-aster complex, 1,100 ± 300 pN s/μm (for the two pronuclei), was about 1/8 of the corresponding estimate for the sea urchin (8,400 pN s/μm)^9^. The difference may be explained again by the size difference. As astral microtubules become longer in the larger sea urchin eggs, it will be harder to move the asters.

Based on past theoretical and experimental considerations using *C. elegans* embryos^8,16–18^, the frictional coefficient of a pronucleus was estimated to be 300 pN s/μm. There is a difference of at least 2-fold between the previous estimate and experimental measurements in the present study (1,100 for the two pronuclei and 640 for one pronucleus [pN s/μm]). The difference may be related to the assumption of a Newtonian fluid in the estimate. In a simple viscous fluid, the frictional coefficient is proportional to the radius of the sphere, and the estimate applies a cytoplasmic viscosity of 0.2 pN s/μm^2^ measured using ∼1 μm beads^16^. In contrast, the real cytoplasm is filled with cytoskeletal filaments and organelles, where a larger object is more difficult to move than a simple object, proportional to the radius. Our measurements indicate that an uncharacterized factor is needed to explain the two-fold or larger difference.

A previous theoretical study predicted that microtubule asters should increase the frictional coefficient 6-fold^8^. An comparison of the speed of sperm- and oocyte-derived pronuclei suggested that the difference is 4.4-fold^18^. The 2-fold difference in the frictional coefficient observed between *zyg-9* (*b244*ts) and wild-type embryos in our study suggests that the effect of microtubule asters is not large, although we must note that *zyg-9* (*b244*ts) does not completely eliminate astral microtubules. Further investigations using *C. elegans* embryos with various mutants and gene knockdowns are needed to characterize the molecular mechanism underlying the force production and frictional coefficient for pronuclear migration.

### Centrosome-organelle mutual pulling model

The mechanisms that produce forces for nuclear centration are still under debate. In most cases, centration depends on microtubule functions. However, it is not clear whether the force for centration is generated by the pushing of microtubules against the cell cortex^14,16,40^ or by the pulling of microtubules by a minus-end directed motor, cytoplasmic dynein^3,41^. The pulling model is further divided into two or more models depending on whether the pulling occurs throughout the cytoplasm^6,9–13,42^ or at the cell cortex^37,39,43^. Pulling at the aster periphery^44^ and at the interphase of aster and actin network^45^ have been proposed recently. The mechanism might differ among species.

The amount of force measured in this study is consistent with the cytoplasmic pulling model and, more specifically, the organelle-centrosome mutual pulling mechanism^3,12,46^. According to the present study, the *C. elegans* embryo generates 14 pN when the nucleus is 1 μm from the center. At this distance, the volume difference of the cytoplasm toward the direction of movement and in the opposite (rear) direction will be ∼5% of the total volume of the embryo. In the organelle-centrosome mutual pulling model, the force toward the center is roughly proportional to the volume difference between the direction of movement and the opposite direction. This means that a force of 14 pN is generated in the 5% volume fraction of the embryo. Thus, we estimate that a total force of ∼300 pN is generated in the cytoplasm if the organelle-centrosome mutual pulling model explains nuclear centration. This is a reasonable amount. Moving single organelles requires several pN of force, indicating that about 100 organelles are moving at a time, which is roughly consistent with the number of moving organelles observed in vivo^12^.

### Combined use of CPM and OI-DIC as a novel tool to characterize forces inside the cell

This study utilized centrifugal force to measure forces inside the cell^47^. We note two major advantages of this novel method. First, it is less invasive than are magnetic or optical tweezers. We confirmed that the embryos hatch after centrifugation, indicating that the procedure does not affect embryogenesis. Furthermore, because it is not necessary to inject large beads into the cell, the experiment can be repeated easily.

Microtubule-based structures responsible for nuclear centration likely remained intact under the centrifugation applied in this study. First, the distance between the nuclei and the center of the cell depended on the centrifugal force (Fig. 3B), consistent with centration activity. Second, the behavior of the nuclei differed between the wild-type cells and *zyg-9-* or *zyg-12*-knockdown cells in which microtubule elongation or associations between microtubules and nuclei are impaired (Fig.5). This result indicates that the microtubules in control cells are not disrupted. Third, after nuclear envelope breakdown, the mitotic spindle rapidly moves back to the center of the cell, even under centrifugation (Supplemental Movies S1 and S2). Because the centering of the spindle is also driven by microtubule-generated forces^16^, microtubule structures for centration of the mitotic spindle remain intact under centrifugation.

Finally, estimation of the stiffness from centrifuge condition (*K*_1_) and that from non-centrifuge condition (*K*_2_) resulted in a similar value (Table 1), supporting that similar centration machinery is working in centrifuge and non-centrifuge condition.

OI-DIC provided a reasonable estimation of the mass density of the cell and its combination with CPM provides a novel method to apply controllable forces to intracellular structures. The application is limited to nuclei currently; however, searching for appropriate biological systems and/or developing protocols to utilize centrifugal forces (e.g., injecting high-density beads into the cell) will provide new ways to investigate the forces acting inside living cells.

## Methods

### C. elegans strains

*C. elegans* strains were cultured using standard procedures^48^. The N2 (wild-type) and XA3507 (*unc-119*(*ed3*)III;*qals3507*[*unc-119*(+) + *Ppie-1*::GFP::LEM-2]III) strains were maintained at 22°C. The DH244 (*zyg-9*(*b244*ts)II) and BW54 (*zyg-12*(*ct350*ts)II) strains were maintained at 16°C, and shifted up to 25°C just before the observation.

### Centrifuge polarizing microscope analyses

The CPM system developed at the Marine Biological Laboratory was used^19^. *C. elegans* embryos were cut out of the adult and embryos before the pseudo-cleavage stage were selected. Embryos together with 4 μL of 0.75× Egg Salt buffer were layered on top of 8 μL of 0.75× Egg Salt buffer with 75% Percoll (vol/vol; Sigma, St. Louis, MO, USA), and spun in the CPM. Differential interference contrast images were obtained using a 40×, 0.55 N.A. objective lens (SLCPlanFI; Olympus, Tokyo, Japan) with a 10×, 0.30 N.A. condenser (UPlanFI; Olympus) with a zoom ocular set to ×1.5 (Nikon, Tokyo, Japan). The specimen was momentarily illuminated by a 532-nm wavelength, 6-ns laser pulse (New Wave Research, Fremont, CA, USA), and images were captured by an interference-fringe-free CCD camera (modified C5948; Hamamatsu Photonics, Hamamatsu, Japan).

For the measurement of the density of the embryo (Fig. 2C), “Density Marker Beads Kit (1.02, 1.04, 1.06, 1.08, 1.09, 1.13g/cc)” (Cospheric, Cat# DMB-kit, Somis, CA, USA) was used. The buffer of the beads were exchanged to 0.75× Egg Salt buffer, and several beads of each density were mounted on the CPM chamber together with *C. elegans* N2 embryos as described above. Images were captured with the rotation speed of 500rpm. The beads of the different density were distinguished by the color.

Time-lapse images were analyzed using ImageJ. The coordinates of the center of the sperm- and oocyte-pronucleus as well as those of the anterior and posterior poles of the cell were quantified by manual tracking. The midpoint of the center of the sperm- and oocyte-pronucleus was defined as the center of the pronuclei. The positions of the centers of the pronuclei along the anterior-posterior (AP) axis after the pronuclear meeting were calculated and plotted against time (Fig. 1D). The plot was fitted to the function *x* = *L*_eq_ × [1 - exp{-(*t* - *t*_0_)/*τ*}] using the Microsoft Excel ‘solver’ tool, where *x* is the position of the centers of the pronuclei (*x* = 0 at cell center, and *x* > 0 for the anterior half), *L*_eq_ is the position where the pronuclei stop moving, *t* is the time, *t*_0_ is the time when the pronuclei pass the center, and *τ* is a characteristic time-scale for the movement. This function is often used to represent trajectories approaching a certain point (*x* = *L*_eq_ in this case) with decreasing speed and fits the experimental data well. At the same time, this formula is a solution of the differential equation *F* - *K x* - *C*_fric_ × (d*x*/d*t*) = 0, where the external force, *F* (e.g., *F*_cfg_), is balanced by a Hookean spring-like centering force, -*Kx*, and a drag force, -*C*_fric_ × (d*x*/d*t*), where *K* is the spring constant and *C*_fric_ is the frictional coefficient. Here, *L*_eq_ = *F*_cfg_/*K* and *τ* = *C*_fric_/*K*. Therefore, the good fit of the experimental data to the formula supports the assumption that the force balance model effectively describes the movement of pronuclei.

### OI-DIC measurement and data analyses

OI-DIC system developed at the Marine Biological Laboratory was used^24,25^. *C. elegans* embryos were cut out of the adult in water and embryos before the pseudo-cleavage stage were selected. Embryos were mounted on a coverslip (pre-coated with 10% poly-L-lysine solution and dried). A coverslip was mounted on a glass slide with a spacer made of VALAP (Vaseline: lanolin: paraffin = 1:1:1) so that the embryo was not compressed. Approximately 30 μL of water was added to fill the space between the coverslip and the glass slide, and the sample was sealed with VALAP. The sample was set on the OI-DIC microscope, equipped with a 40×/0.60 N.A. objective lens (LUCPlanFLN; Olympus) and a yellow 576 nm bandpass filter (FF01-576/10-25, Semrock, Rochester, NY, USA). Images were recorded with a CCD camera (Teledyne Lumenera, Infinity3-1M, Ottawa, Ontario, Canada) and processed to calculate the optical path difference (OPD) as described^24,25,49,50^.

The obtained images with OPD values in each pixel were analyzed using ImageJ. A line with 10 pixels that passes through the center of a nucleus and perpendicular to the long axis of the ellipsoidal embryo was drawn by hand. The OPD along the line was quantified using the ‘Plot Profile’ function. OPD is defined as the difference in refractive index (ΔRI) multiplied by thickness of the specimen^29^. The OPD values along the line (*x*-axis) were considered as the superposition of the following three functions.

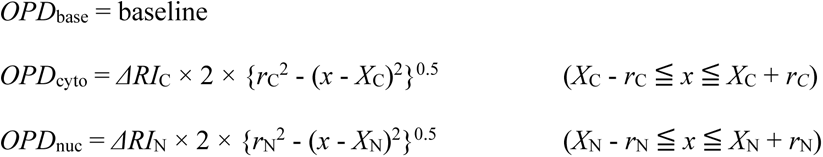

Here, *OPD*_base_, *OPD*_cyto_, and *OPD*_nuc_ are the contributions to the OPD of the background (water), cytoplasm, and nucleus, respectively. *ΔRI*_C_ and *ΔRI*_N_ are the difference in the refractive index of the cytoplasm and nucleus compared with that of water, *r*_C_ and *r*_N_ are the radii of the cytoplasm (cross section) and nucleus, and *X*_C_ and *X*_N_ are the center coordinates along the *x*-axis of the cytoplasm and nucleus. The OPD profile was fitted to the superposition of the above functions using the Microsoft Excel ‘solver’ tool. The obtained difference in refractive indices compared with that of water (*ΔRI*_C_ and *ΔRI*_N_) were converted to the concentration of dry mass (*C*_dm_, in g/mL) using *C*_dm_ = *ΔRI* / *α*, where *α* is 0.0018 (100 mL/g)^29^. Finally, the density of the cytoplasm or nucleus (*D*_C_ or *D*_N_, in kg/m^3^) can be calculated according to the following formula.

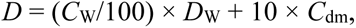

where *C*_W_ is the percentage of water in the cytoplasm or nucleus (% or g/100 mL), *D*_W_ is the density of water (997 kg/m^3^ at 25°C), and 10 × *C*_dm_ is the dry mass in kg/m^3^. *C*_W_ can be determined as *C*_W_ = (100 – *V*_sp_ × *C*_dm_), assuming that the specific volume of the cytoplasm and the nucleoplasm is the same as the specific volume of proteins (*V*_sp_ = 0.75 mL/g)^29^.

### Measurements of the nuclear volume using conventional confocal microscopes

To quantify the nuclear volume (*NV*), images of embryos of the strain XA3507 (*unc-119*(*ed3*)III;*qals3507*[*unc-119*(+) + P*pie-1*::GFP::LEM-2]III) in which the nuclear membrane was fluorescently labeled were obtained. Embryos were placed in 0.75× egg-salt and images were obtained at room temperature (22–24°C) using a spinning-disk confocal system (CSU-X1; Yokogawa, Tokyo, Japan) mounted on an inverted microscope (IX71; Olympus) equipped with a 60×, 1.30 N.A. objective (UPLSAPO 60XS2; Olympus). Images were acquired with a CCD camera (iXon; Andor Technology, Belfast, UK) controlled using Metamorph (version 7.7.10.0). Images were acquired with a z-interval of 1 μm, and the outline of the pronuclei was traced by hand using Imaris (Oxford Instruments, Abingdon, UK). The surfaces of the pronuclei were reconstituted and the volumes were calculated using Imaris.

### Measurements of the nuclear migration in non-centrifuge condition

The speed of pronuclear migration in the non-centrifuge condition was quantified as follows. Images of the wild-type N2 strain were obtained using a Nomarski DIC microscope (Olympus BX51) with a 100× oil immersion objective lens at a 2-s time interval. The 2-dimentional coordinates of the center of the sperm-derived pronucleus was quantified manually using ImageJ. Because migration occurred mainly along the long axis of the embryo, the 2-dimensional positions of the pronuclei were projected onto the long axis. The position(*x*)-vs-time(*t*) plot (Fig. 4A) was fitted to the formula, *x* = *X*_0_ exp{-*R*_KC_ (*t* - *t*_0_)}, using the Microsoft Excel ‘solver’ tool. For the fitting, we used the data from when the position exceeds the halfway after the meeting until the position reaches the cell center, or the maximum value (Fig. 4A, filled circle).

### Evaluation of combined errors

The variance (error) of the calculated values was evaluated by the law of propagation of uncertainty. With this law, if the parameter *y* is a function of *x*_i_ (i.e., *y* = f(*x*_1_, *x*_2_, …) the variance of *y* (σ^2^(*y*)) is calculated as σ^2^(*y*) = Σ_i_{(∂f/∂*x*_i_)^2^×σ^2^(*x*_i_)}. In this study, the variances of *K*_1_, *C*_fric_, and *K*_2_ were evaluated as follows. See Table 1 for definitions of the symbols.

*F*_cfg_ = *NV* × Δ*ρ* × RCF×*g*

*K*_1_ = *F*_cfg_ / *L* = *NV*×Δ*ρ*×*g*×RCF/*L*_eq_ = *NV*×Δ*ρ*×*g*/*β*_RL_

*σ*^2^(*K*_1_) = (*NV*^2^ *g*^2^/*β* ^2^)×*σ*^2^(Δ*ρ*) + (Δ*ρ*^2^ *g*^2^/*β* ^2^)×*σ*^2^(*NV*) + (*NV*^2^Δ*ρ*^2^ *g*^2^/*β* ^4^)×*σ*^2^(*β*_RL_)

*C*_fric_ = *NV*×Δ*ρ*×*g*/*β*_RVc_.

*σ*^2^(*C*_fric_) = (*NV*^2^ *g*^2^/*β*_RVc_^2^)×*σ*^2^(Δ*ρ*) + (Δ*ρ*^2^ *g*^2^/*β*_RVc_^2^)×*σ*^2^(*NV*) + (*NV*^2^Δ*ρ*^2^ *g*^2^/*β*_RVc_^4^)×*σ*^2^(*β*_RVc_)

*K*_2_ = *C*_fric_ × *R*_KC_.

*σ*^2^(*K*_2_) = (*R*_KC_^2^)×*σ*^2^(*C*_fric_) + (*C*_fric_^2^)×*σ*^2^(*R*_KC_)

## Acknowledgements

We thank the late Shinya Inoué for inventing the CPM and for discussions about its use. We also thank Shinya Inoué and Marine Biological Laboratory for letting us use the CPM and later letting us transfer the CPM to the National Institute of Genetics, Japan. We thank Tomoko Ozawa, Yuki Hara, Tomo Kondo, Kazunori Yamamoto, and Kenji Kimura for technical assistance. We thank Hajime Takahashi (Olympus) for his support in moving the CPM to Japan. We thank Hirokazu Tanimoto, Yuta Shimamoto, Saya Ichihara, Ken Fujii, Tokitaka Katayama for their valuable comments on the manuscript, and the members of the Kimura lab for discussion. This work was supported by a Whitman Center Fellowship from the Marine Biological Laboratory (to AK) and by the JSPS KAKENHI (grant numbers JP18H02414 and JP18H05529 to AK, and JP18KK0202 to AK and GG). MS gratefully acknowledges funding from the NIGMS/NIH under Grant Nr. R01GM101701 and MS and RO from the Marine Biological Laboratory through the Inoué Endowment Fund.

