## Supplemental Figure S1, Movie legends for "Live-cell imaging under centrifugation characterized the cellular force for nuclear centration in the *Caenorhabditis elegans* embryo"

[6] Independent Researcher

[7] Biomedical Research Institute, National Institute of Advanced Industrial Science and Technology, Ikeda, Osaka, Japan.

*Corresponding author

**Supplemental Figure S1. Quantification of the nuclear volume.**

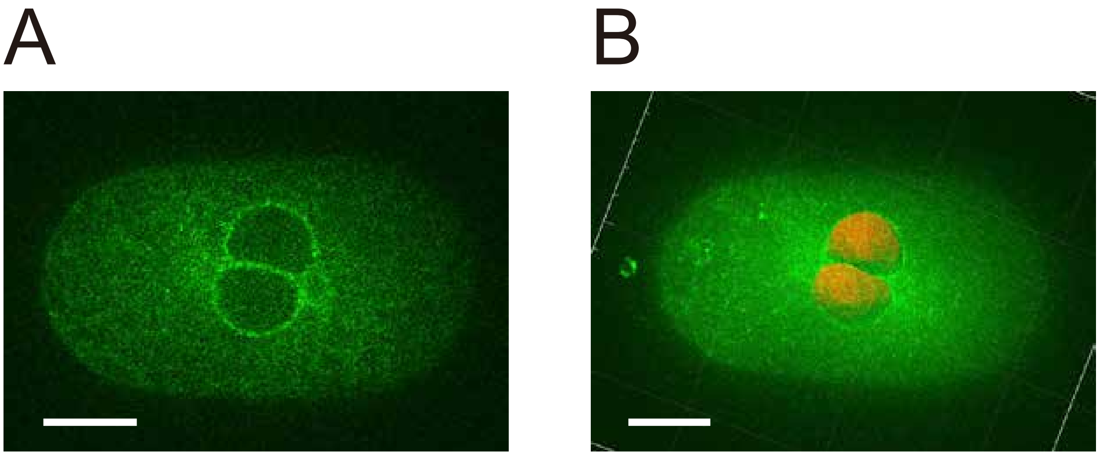


(A) Representative confocal image of the *C. elegans* embryo in the pronuclear migration stage. The nuclear membrane is visualized with the green fluorescence protein. (B) Manual segmentation of the nuclear membrane from the image in (A) to quantify the nuclear volume. The 3-dimensional image with nuclear surface shown in red was drawn using Imaris. Scale bar, 10 μm.

**Supplemental Movie S1.** Time-lapse movie of the *C. elegans* embryo mounted in the CPM with a rotation speed of 500 rpm (20 ×*g*, corresponding to Fig. 1B). Play back speed ×100.

**Supplemental Movie S2.** Time-lapse movie of the *C. elegans* embryo mounted in the CPM with a rotation speed of 3,000 rpm (780×*g*, corresponding to Fig. 1C and Fig. 1D(ii)). Initially, the rotation speed was kept 500 rpm (20 ×*g*) until just before the meeting (“4:56:57” at the second from left at the bottom), and then the speed was increased. The speed reached 3,000 rpm (780 ×*g*) 50 s later (“4:57:47”). Play back speed ×100.
